# *Cryptococcus neoformans* Chitin Synthase 3 (Chs3) Plays a Critical Role in Dampening Host Inflammatory Responses

**DOI:** 10.1101/814400

**Authors:** Camaron R. Hole, Woei C. Lam, Rajendra Upadhya, Jennifer K. Lodge

## Abstract

*Cryptococcus neoformans* infections are significant causes of morbidity and mortality among AIDS patients and the third most common invasive fungal infection in organ transplant recipients. One of the main interfaces between the fungus and the host is the fungal cell wall. The cryptococcal cell wall is unusual among human pathogenic fungi in that the chitin is predominantly deacetylated to chitosan. Chitosan deficient strains of *C. neoformans* were found to be avirulent and rapidly cleared from the murine lung. Moreover, infection with a chitosan deficient *C. neoformans* lacking three chitin deacetylases (*cda1Δ2Δ3Δ*) was found to confer protective immunity to a subsequent challenge with a virulent wild type counterpart. In addition to the chitin deacetylases, it was previously shown that chitin synthase 3 (Chs3) is also essential for chitin deacetylase mediated formation of chitosan. Mice inoculated with *chs3Δ* at a dose previously shown to induce protection with cda*1Δ2Δ3Δ* die within 36 hours after installation of the organism. Mortality was not dependent on viable fungi as mice inoculated with heat-killed preparation of *chs3Δ* died at the same rate as mice inoculated with live *chs3Δ*, suggesting the rapid onset of death was host mediated likely caused by an over exuberant immune response. Histology, cytokine profiling, and flow cytometry indicates a massive neutrophil influx in the mice inoculated with *chs3Δ*. Mice depleted of neutrophils survived *chs3Δ* inoculation indicating that death was neutrophil mediated. Altogether, these studies lead us to conclude that Chs3, along with chitosan, plays critical roles in dampening cryptococcal induced host inflammatory responses.

**IMPORTANCE:** *Cryptococcus neoformans* is the most common disseminated fungal pathogen in AIDS patients, resulting in ∼200,000 deaths each year. There is a pressing need for new treatments for this infection, as current antifungal therapy is hampered by toxicity and/or the inability of the host’s immune system to aid in resolution of the disease. An ideal target for new therapies is the fungal cell wall. The cryptococcal cell wall is different than many other pathogenic fungi in that it contains chitosan. Strains that have decreased chitosan are less pathogenic and strains that are deficient in chitosan are avirulent and can induce protective responses. In this study we investigated the host responses to *chs3*Δ, a chitosan-deficient strain, and found mice inoculated with *chs3*Δ all died within 36 hours and death was associated with an aberrant hyperinflammatory immune response driven by neutrophils, indicating that chitosan is critical in modulating the immune response to *Cryptococcus*.

## Introduction

*Cryptococcus neoformans* is a ubiquitous encapsulated fungal pathogen that causes pneumonia and meningitis in immunocompromised individuals. *C. neoformans* is the most common disseminated fungal pathogen in AIDS patients, with an estimated quarter million cases of cryptococcal meningitis each year resulting in ∼200,000 deaths (1, 2) and remains the third most common invasive fungal infection in organ transplant recipients (3). Current antifungal therapy is often hampered by toxicity and/or the inability of the host’s immune system to aid in resolution of the disease; treatment is further limited by drug cost and availability in the resource-limited settings (4). The acute mortality rate of patients with cryptococcal meningitis is between 10-30% in medically-advanced countries (5, 6), and even with appropriate therapy at least one third of patients with cryptococcal meningitis will undergo mycologic and/or clinical failure (4). Patients that do recover can be left with profound neurological sequelae, highlighting the need for more effective therapies, and/or vaccines to combat cryptococcosis

One of the main interfaces between the fungus and the host is the fungal cell wall. Most fungal cell walls contain chitin, however, the cryptococcal cell wall is unusual in that the chitin is predominantly deacetylated to chitosan. Chitin is a homopolymer of β-1,4-linked *N*-acetylglucosamine (GlcNAc) and is one of the most abundant polymers in nature. Immunologically, chitin can induce allergy and strong Th2-type immune responses (7). Chitin is polymerized from cytoplasmic pools of UDP-GlcNAc by a multiple trans-membrane protein chitin synthase (CHS) and there are eight Chs’s encoded in *C. neoformans* genome (8). Chitosan, the deacetylated form of chitin, is generally less abundant in nature than chitin, but is found in the cell wall of several fungal species depending on growth phase (8). Chitosan is not synthesized de novo but is generated from chitin through enzymatic conversion of GlcNAc to glucosamine by chitin deacetylases (CDAs) and *C. neoformans* makes three CDAs (9). Why *Cryptococcus* converts chitin to chitosan and what advantages this conversion provides to the organism are not well understood.

Deletion of a specific chitin synthase (CHS3), or deletion of all three chitin deacetylases causes a significant reduction in chitosan in the vegetative cell wall (9). These chitosan deficient strains of *C. neoformans* are avirulent and rapidly cleared from the murine lung (9). Moreover, infection with a chitosan deficient *C. neoformans* strain lacking three chitin deacetylases (*cda1Δ2Δ3Δ*) was found to confer protective immunity to a subsequent challenge with a virulent wild type counterpart (10). These findings suggest that there is an altered host response to chitosan-deficient strains. Therefore, we wanted to determine the nature of host immune response to an infection with chitosan deficiency caused by the deletion of *C. neoformans CHS3* gene.

Surprisingly, we observed that all mice inoculated with chitosan-deficient *chs3*Δ died within 36 hours. Death was not dependent on live organism or the mouse background. We hypothesized that the rapid onset of mortality was likely due to an aberrant immune response. Histology, cytokine profiling, and flow cytometry indicates a massive influx of neutrophils in the mice inoculated with *chs3*Δ. Mice depleted of neutrophils all survived inoculation of the *chs3*Δ strain, indicating that the observed mortality is neutrophil mediated. Together, these results suggest that chitin synthase 3 is important in modulating the immune response to *Cryptococcus*.

## Results

### Complete deletion and complementation of C. neoformans chitin synthase 3 (Chs3)

With better annotation of the cryptococcal genome, we found that our previously reported *chs3*Δ strain (8) was not a complete deletion. While the protein is not functional as the catalytic domain was deleted, the original strain still harbored 689 bp of gene sequence potentially sufficient to encode a ∼25 kDa protein. As this gene is highly expressed under vegetative growth, the truncated protein might influence either the virulence or the nature of the host immune response to the mutant. Due to this, we generated a complete deletion of the chs3 gene, including the 5’ UTR to delete the promoter as well, in KN99 by biolistic transformation. All the isolates were characterized by diagnostic PCR screening and southern blot hybridization.

The original *chs3*Δ strain exhibited a large number of phenotypes including changes in morphology including 2-3 fold enlarged cells and a budding defect, temperature sensitivity, leaky melanin, and chitosan deficiency, among others (8). Cells of the new *chs3*Δ strain exhibited the same morphologic changes observed in the original strain (Fig. 1A). Additionally, the new *chs3*Δ strain is also temperature sensitive (Fig. 1B) and deficient in chitosan (Fig. 1C).

**Figure 1.**
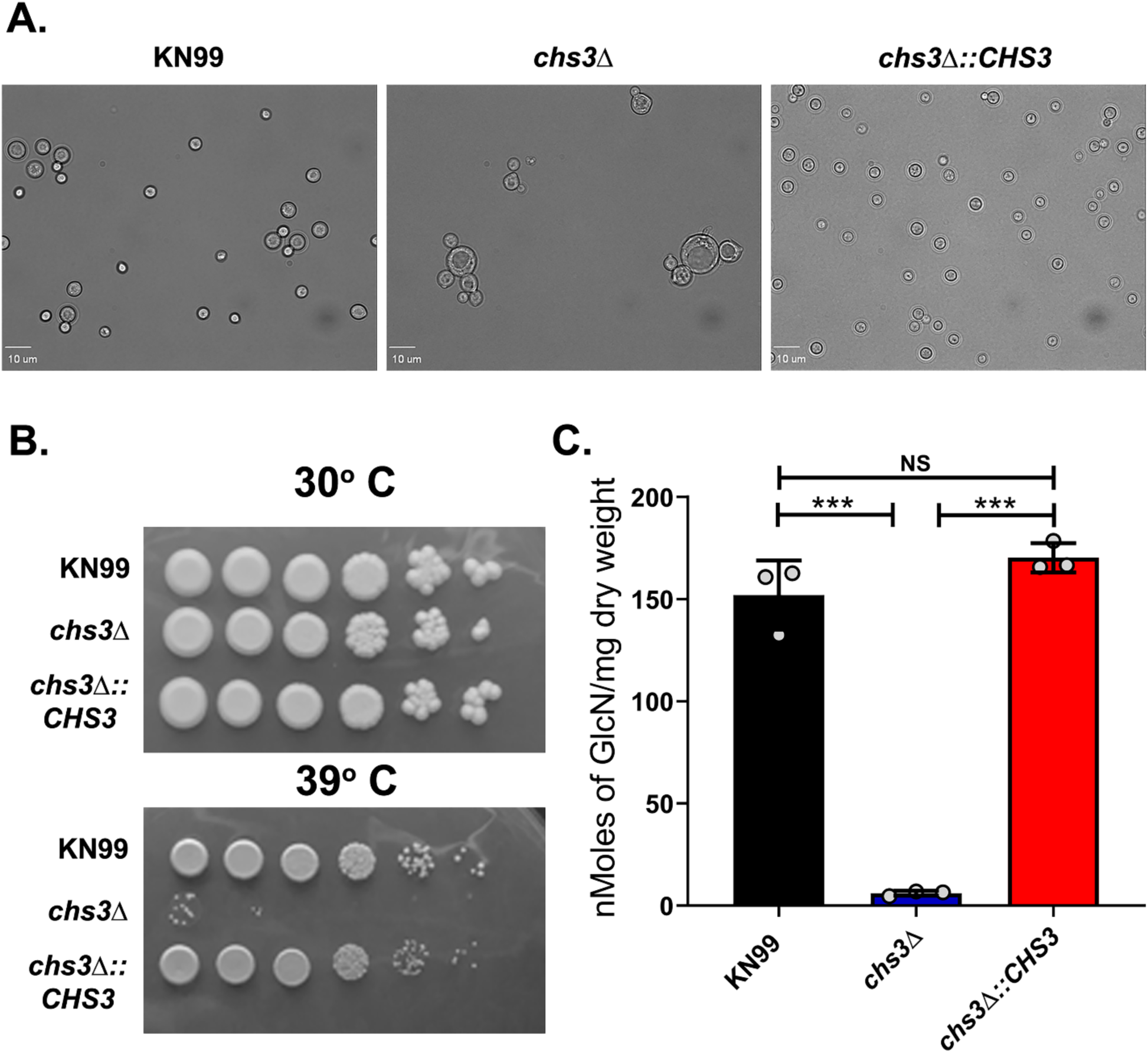
Deletion and complementation of *C. neoformans* chitin synthase 3 (Chs3): (A) For morphological analysis, cells were incubated for 2 days in YPD and diluted to an OD_650_ of 0.2 with PBS. Five microliters each cell solution was spotted on to a clean glass slide and photographed at 40X. (B) Temperature sensitivity. Cultures were grown overnight in YPD then diluted to an OD650 of 1.0. Tenfold serial dilutions were made in PBS and 3µl of each was plated. The plates were grown for 4 days at 30°C or 39°C. (C) Quantitative determination of cell wall chitosan by the MBTH assay. Cells were grown in YPD for two days, collected, washed and used for the assay. Data represents the average of three biological experiments ± standard deviation (SD) and are expressed as nMoles of glucosamine per mg dry weight of yeast cells (***, *P* < 0.001).

Previously we attempted to complement the original *chs3*Δ strain a multitude of ways and all attempts failed, leading us to conclude that the cell wall of the *chs3*Δ strain was compromised to a point that they could not survive any of the transformation procedures. (9). With this in mind, we attempted to complement the new *chs3*Δ strain using electroporation into the endogenous locus, replacing the NAT resistance marker and were successful. The complemented strain (*chs3Δ::CHS3)* reversed all the observed phenotypes including the changes in morphology, temperature sensitivity and chitosan deficiency (Fig. 1A-C).

### Inoculation with the *chs3*Δ strain induces rapid mouse mortality

We have previously shown that chitosan is essential for growth in the mammalian host. Strains with three different chitosan deficiency genotypes (*chs3*Δ, *csr2Δ*, and *cda1Δ2Δ3Δ*) all show rapid pulmonary clearance in a mouse model of cryptococcosis and complete loss of virulence (9). Mice that received a high inoculation (10^7^ CFU) of the chitosan deficient strain *cda1Δcda2Δcda3Δ* were able to clear the infection and were found to be protected against a subsequent challenge with wild-type KN99 (WT) *C. neoformans* (10). Notably, this chitosan deficient strain is protective even when heat-killed (10). Protective immunization is dependent on the inoculum size, as only mice that received 10^7^ CFU of *cda1Δcda2Δcda3Δ* were protectively immunized, mice that received a lower inoculation were not protected (10).

Based on these data, we set out to test whether inoculation with other chitosan deficient strains would also confer protection. We started this process using the new *chs3*Δ strain which is chitosan deficient (Fig. 1C). We inoculated C57BL/6 mice intranasally with 10^7^ CFU of live *cda1Δ2Δ3Δ* (a concentration that is shown to be protective for *cda1Δ2Δ3Δ*)*, chs3Δ, chs3Δ::CHS3*, or WT *C. neoformans* KN99 and were monitored for survival. As expected, mice that received *cda1Δ2Δ3Δ* all survived the infection and mice that received the WT KN99 or *chs3Δ::CHS3* all died or were euthanized due to morbidity around day 6 (KN99) or day 8 (*chs3Δ::CHS3)* post inoculation with this high inoculum (Fig. 2). What was surprising, however, was that the mice inoculated with *chs3*Δ all died within 36 hours after instillation of the organism (Fig. 2).

**Figure 2.**
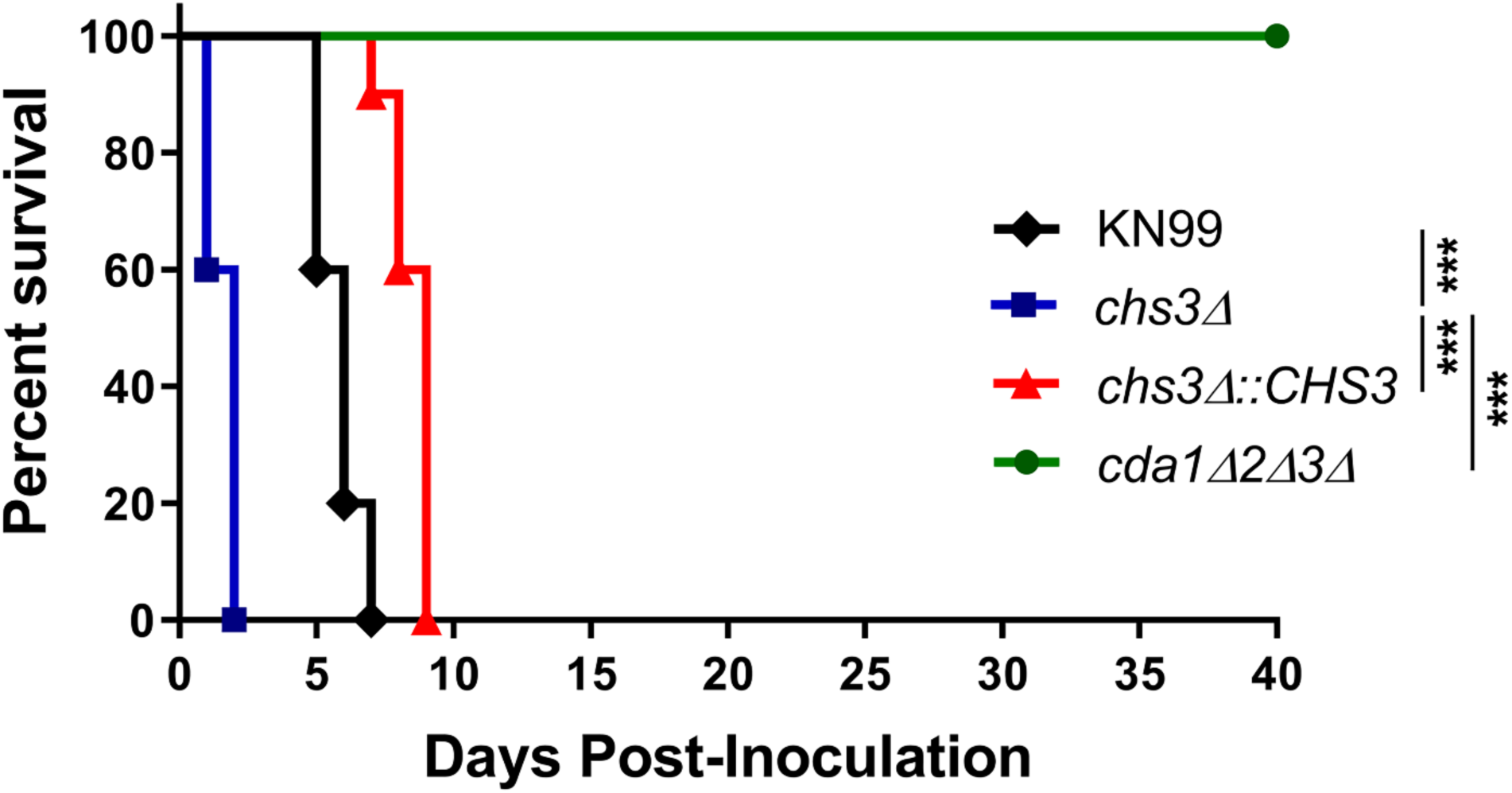
Inoculation with the *chs3*Δ strain induces rapid mouse mortality: C57BL/6 mice were infected with 10^7^ live CFUs of each strain by intranasal inoculation. Survival of the animals was recorded as mortality of mice for 40 days post inoculation. Mice that lost 20% of the body weight at the time of inoculation or displayed signs of morbidity were considered ill and sacrificed. Data is representative of one experiment with 5 mice for KN99, 5 mice for *cda1Δ2Δ3Δ*, 10 mice for *chs3Δ*, and 10 mice for *chs3Δ::CHS3*. Virulence was determined using Mantel-Cox curve comparison with statistical significance determined by log-rank test. (***, *P* < 0.001).

The rapid rate of mortality suggested death was not due to fungal proliferation or burden. Furthermore, we previously showed that the original *chs3*Δ strain is rapidly cleared from the host at a lower inoculum (9). Based on these finding, we tested if mortality was dependent on viable fungi. We heat-killed (HK) WT KN99, *chs3Δ*, and *chs3Δ::CHS3* strains at 70°C for 15 minutes. Complete killing was confirmed by plating for CFUs. C57BL/6 mice then received an intranasal inoculation with 10^7^ CFU of HK WT KN99, HK *chs3*Δ or HK *chs3Δ::CHS3* and were monitored for survival. Mice that received HK WT KN99 or HK *chs3Δ::CHS3* all survived the inoculation of heat killed cells (Fig. 3A). Conversely, mice that received HK *chs3*Δ all died at the same rate as observed above with live *chs3*Δ (Figs. 2 and 3A) indicating that mortality was not dependent on the viability of the fungi. In addition, to confirm that the observed phenotype was due to loss of Chs3 and not some secondary mutation in the strain used, we assayed the original *chs3*Δ mutant strain (Supplemental figure 1) and saw the same rapid mortality observed in Figure 3A.

**Figure 3.**
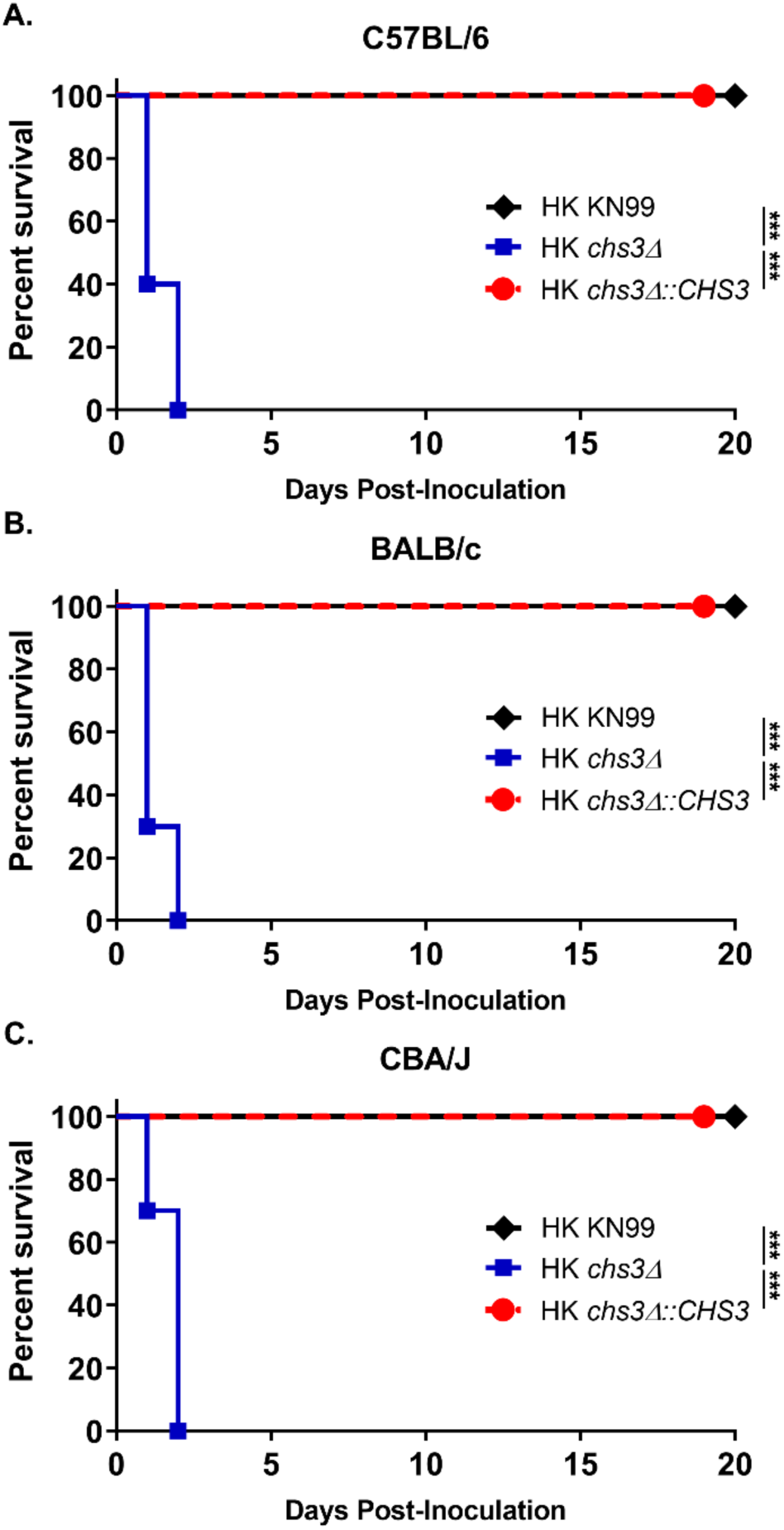
Mortality is not dependent on the viability of the fungi or mouse background: (A) C57BL/6, (B) BALB/c, or (C) CBA/J mice were inoculated with 10^7^ Heat-killed CFUs of each strain by intranasal inoculation. Survival of the animals was recorded as mortality of mice for 20 days post inoculation. Mice that lost 20% of the body weight at the time of inoculation or displayed signs of morbidity were considered ill and sacrificed. Data is cumulative of one experiment with 5 mice for KN99, and two experiments with 5 mice for *chs3*Δ and *chs3Δ::CHS3* each for a total of 10 mice. Virulence was determined using Mantel-Cox curve comparison with statistical significance determined by log-rank test. (***, *P* < 0.001).

Different mouse backgrounds have various susceptibilities to *C. neoformans* depending on the strain used (11). Due to the strong phenotype observed with *chs3*Δ in the C57BL/6 mice, we wanted to verify that rapid rate of mortality was not due to the mouse background. To assess susceptibility in different mouse backgrounds, BALB/c or CBA/J mice received an intranasal inoculation with 10^7^ CFU of HK WT KN99, HK *chs3*Δ or HK *chs3Δ::CHS3* and were monitored for survival. Regardless of the mouse background, mice that received HK WT KN99 or HK *chs3Δ::CHS3* all survived the challenge, whereas mice that received HK *chs3*Δ all died at the same rate as observed in the C57BL/6 mice (Figs. 3A-C) indicating that rapid rate of mortality was not a mouse background phenomenon.

### A massive inflammatory response is triggered to *chs3*Δ inoculation

The above data indicates that mice are not dying due to the fungal burden, as death was not dependent on viable fungi in multiple mouse backgrounds (Figure 3). These data suggest that the mortality associated with *chs3*Δ maybe host mediated (12). To test this, C57BL/6 mice received an intranasal inoculation with 10^7^ CFU HK WT KN99, HK *chs3*Δ or HK *chs3Δ::CHS3* and the lungs were processed for histology. For all immune studies, we chose to use heat-killed fungi to control for fungal burden as the WT KN99 and *chs3Δ::CHS3* would rapidly outgrow the *chs3*Δ strain and potentially skew our results. Lungs were processed at 8 hours post inoculation as we could not constantly keep the *chs3*Δ inoculated mice alive for the full 24h. The 8-hour time point was chosen as this was the time the animal started to show signs of morbidity. The paraffin embedded lungs were sectioned and processed for hematoxylin-eosin (H&E) staining. Histological analysis of the infected lung show little pathology in the lung of the mice inoculated with either HK KN99 or HK *chs3Δ::CHS3* compared to the strong inflammatory response in the lungs of the *chs3*Δ inoculated mice at 8h (Figure 4). The lungs from mice inoculated with *chs3*Δ exhibit abundant foci of inflammation spread across the whole lung section (Fig 4C) consisting of a profound amount of mixed inflammatory infiltrates with enhanced presences of granulocytes (Figs 4D-E). Such severe pneumonia and lung damage could explain the mortality observed in *chs3*Δ inoculated mice and indicates that the immune response in the lungs, albeit robust, is nonprotective and detrimental.

**Figure 4.**
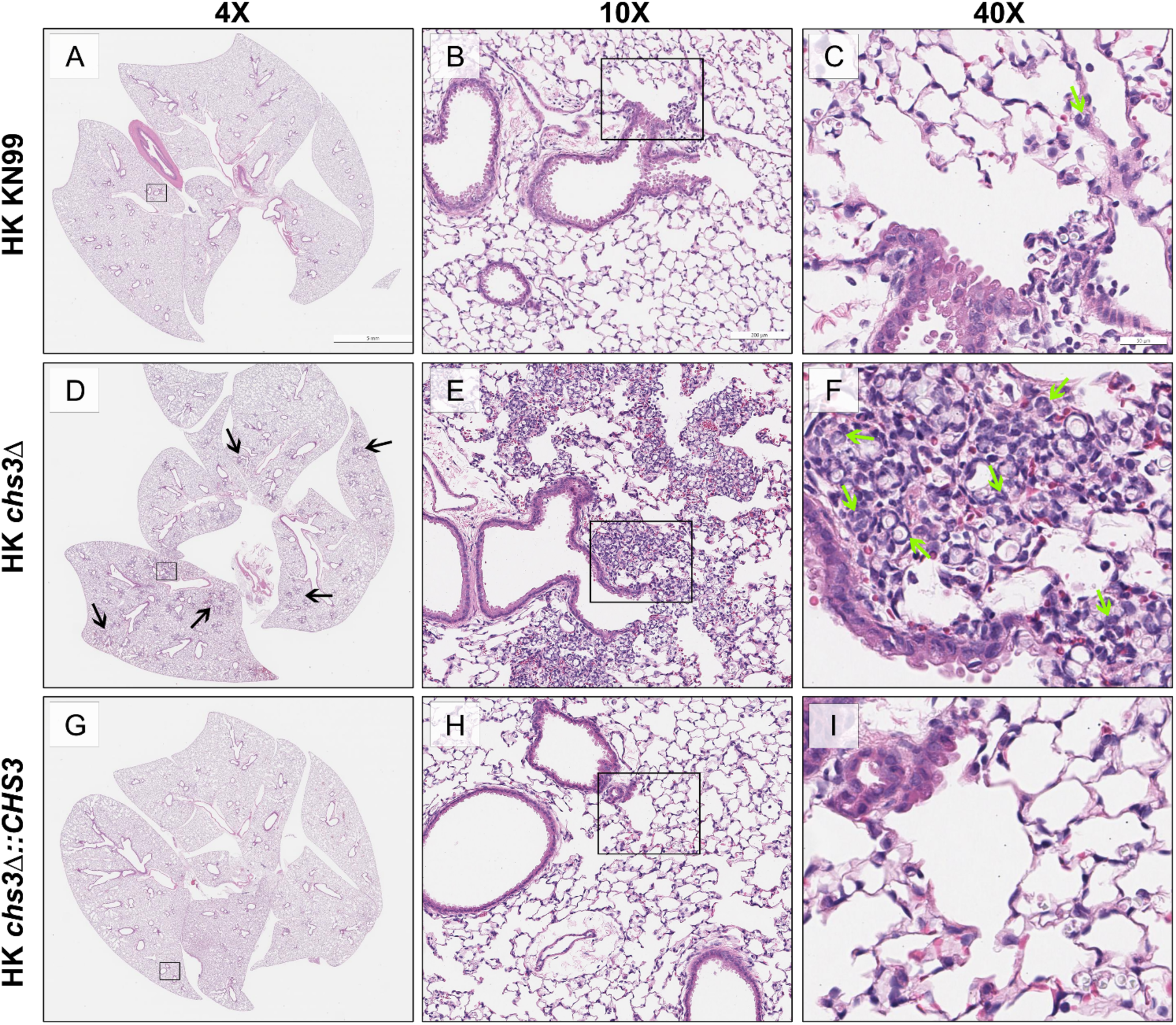
Massive inflammatory response is triggered to *chs3Δ:* C57BL/6 mice were inoculated with 10^7^ Heat-killed CFUs of each strain by intranasal inoculation. At 8 hours post inoculation, the lungs were harvested, embedded, sectioned and processed for hematoxylin-eosin staining. HK KN99 inoculated mice (A-C) display a limited inflammatory response. HK *chs3*Δ inoculated mice (D-F) exhibit abundant foci of inflammation (black arrows) spread across the whole lung section consisting of a profound amount of mixed inflammatory infiltrates with enhanced presences of neutrophils (green arrows). HK *chs3Δ::CHS3* inoculated mice (G-I) displayed a similar limited inflammatory response observed in the HK KN99 inoculated mice. Images are representative images of two independent experiments using three mice per group.

### *chs3*Δ induces a strong proinflammatory cytokine response

Because we observed massive infiltration of immune cells in the lungs of *chs3*Δ inoculated mice (Figs 4D-E), we next assessed the cytokine/chemokine produced. To do this, C57BL/6 mice received an intranasal inoculation with 10^7^ CFU HK WT KN99, HK *chs3*Δ or HK *chs3Δ::CHS3* and at 8hr post inoculation homogenates were prepared from the lungs of each group as well as a PBS control group. Cytokine/chemokine responses were determined from the lung homogenates using the Bio-Plex Protein Array System. We observed an increase in multiple cytokines (Supplementary Figure 2), however there was a significant increase in the chemokines KC (Fig. 5A) and G-CSF (Fig. 5B), as well as extremally high levels of IL-6 (Fig. 5C) in *chs3*Δ inoculated mice compared to PBS, HK WT KN99 or HK *chs3Δ::CHS3* inoculated mice. This cytokine profile is indicative of a strong neutrophilic response in the lungs which correlates with the histology data above indicating an enhanced presence of granulocytes (Figure 4).

**Figure 5.**
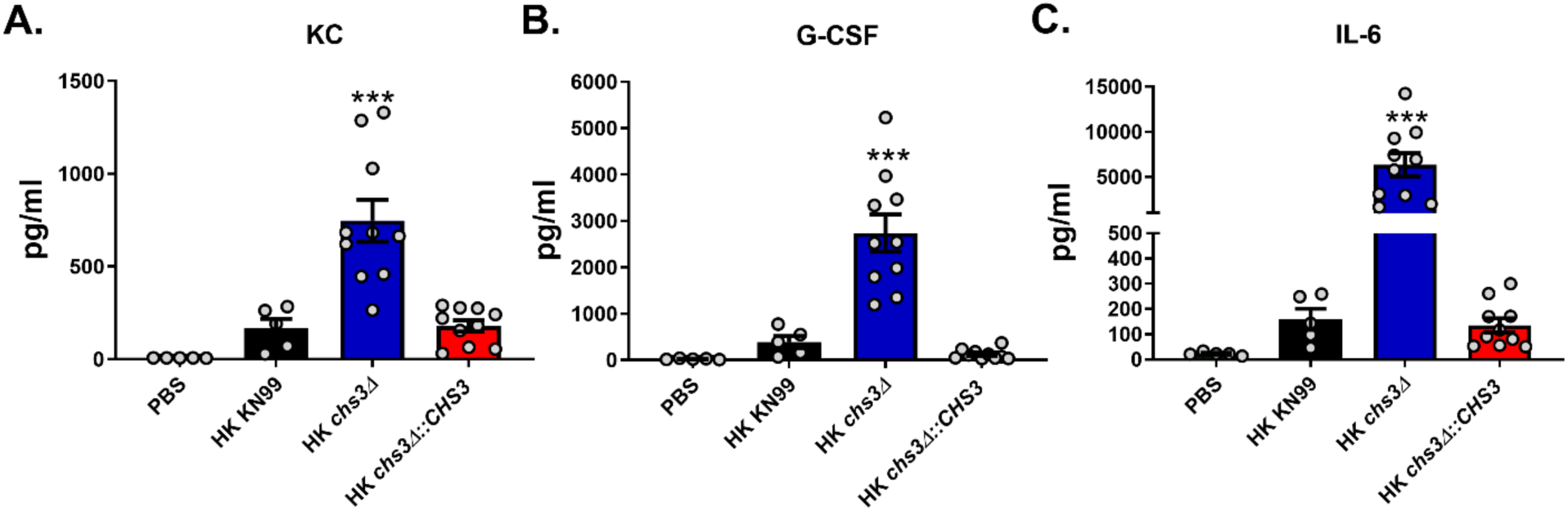
*chs3*Δ induces a strong proinflammatory cytokine response: C57BL/6 mice were inoculated with 10^7^ Heat-killed CFUs of each strain by intranasal inoculation. At 8 hours post inoculation, homogenates were prepared from the lungs of each group. Cytokine/chemokine responses were determined from the lung homogenates using the Bio-Plex Protein Array System. Data is cumulative of one experiment with 5 mice for PBS and KN99, and two experiments with 5 mice for *chs3*Δ and *chs3Δ::CHS3* each for a total of 10 mice experiments, ± standard errors of the means (SEM). Each dot represents data from an individual mouse. (***, *P* < 0.001)

### A significant increase in neutrophil recruitment in the lungs of *chs3*Δ inoculated mice

Since both the histology and cytokine analysis indicates a strong inflammatory response, we wanted to identify the responding cells. For this, C57BL/6 mice received an intranasal inoculation with 10^7^ CFU of HK WT KN99, HK *chs3*Δ or HK *chs3Δ::CHS3* and at 8hr post inoculation pulmonary leukocytes were isolated from the lungs of each group by enzymatic digestion and subjected to flow cytometry analysis for leukocyte identity (Supplementary Figure 3). Consistent with the above histology data, there was a significant increase in the total number of immune cells in the lungs of *chs3*Δ inoculated mice (Fig. 6A). In addition, there was a significant increase in both the total number and percent neutrophils in the lungs *chs3*Δ inoculated mice compared to the WT KN99 or HK *chs3Δ::CHS3* inoculated mice (Figs 6B-C). We did not observe a significant change in any of the other cell types assayed (Supplementary Figure 4).

**Figure 6.**
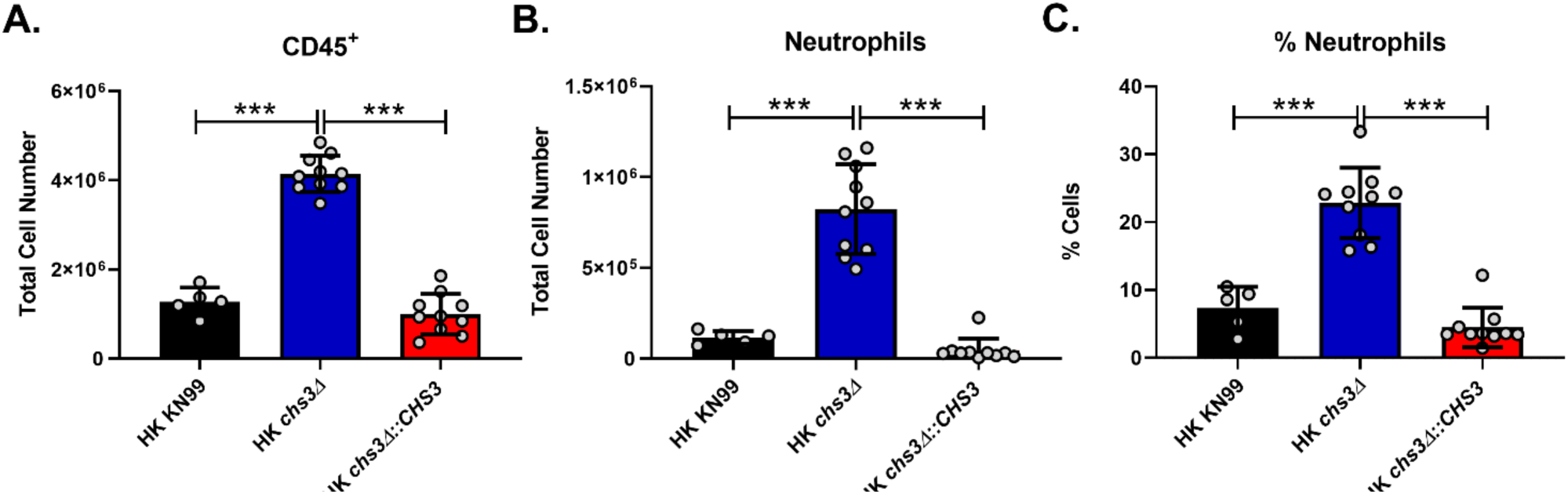
A significant increase in neutrophil recruitment in *chs3*Δ inoculated mice. C57BL/6 mice were inoculated with 10^7^ Heat-killed CFUs of each strain by intranasal inoculation. At 8 hours post inoculation, pulmonary leukocytes were isolated from the lungs of each group and subjected to flow cytometry analysis (see Supplemental Table 2 for antibodies and Supplemental Fig. 3 for gating strategy). (A) total cell number of leukocytes. (B) Total and (C) percent neutrophils (CD11b^+^/CD24^+^/Ly6G^+^/CD45^+^). Data is cumulative of one experiment with 5 mice for PBS and KN99, and two experiments with 5 mice for *chs3*Δ and *chs3Δ::CHS3* each for a total of 10 mice experiments, ± standard errors of the means (SEM). Each dot represents data from an individual mouse. (***, *P* < 0.001)

### Depletion of neutrophils protects *chs3*Δ inoculated mice

Due to the significant increase in neutrophil recruitment to the lungs of mice inoculated with the *chs3*Δ strain, we sought to determine the role of neutrophils in the rapid mortality observed in these animals. To test this, C57BL/6 mice were injected with 200ug of anti-Ly6G (1A8), an antibody that specifically depletes neutrophils (13, 14)) or an isotype antibody 24 hours before intranasal inoculation with 10^7^ CFU of HK *chs3*Δ and monitored for survival. Mice were injected with antibody every 24 hours for the first 5 days post challenge. After day 5, the mice were injected every 48 hours. This antibody is usually injected every 48 hours, however, with the high number of neutrophils recruited (Fig. 6) and the elevated levels of neutrophil growth factors (Fig. 5) we elected to increase the number of the initial injections to ensure neutrophil depletion. Mice treated with the isotype antibody all died at the same rate as observed above with HK *chs3*Δ (Figs. 3A and 7A), whereas mice that were treated with anti-Ly6G all survived (Fig. 7A) indicating that death was neutrophil mediated. To confirm this finding, we repeated the experiment in BALB/c and CBA/J mice. Consistent with our findings in C57BL/6, mice that were depleted of neutrophils all survived inoculation with HK *chs3*Δ whereas mice treated with the isotype antibody all died regardless of mouse background (Figs 7B-C). These data demonstrate that the rapid rate of mortality observed in mice inoculated *chs3*Δ is neutrophil dependent.

**Figure 7.**
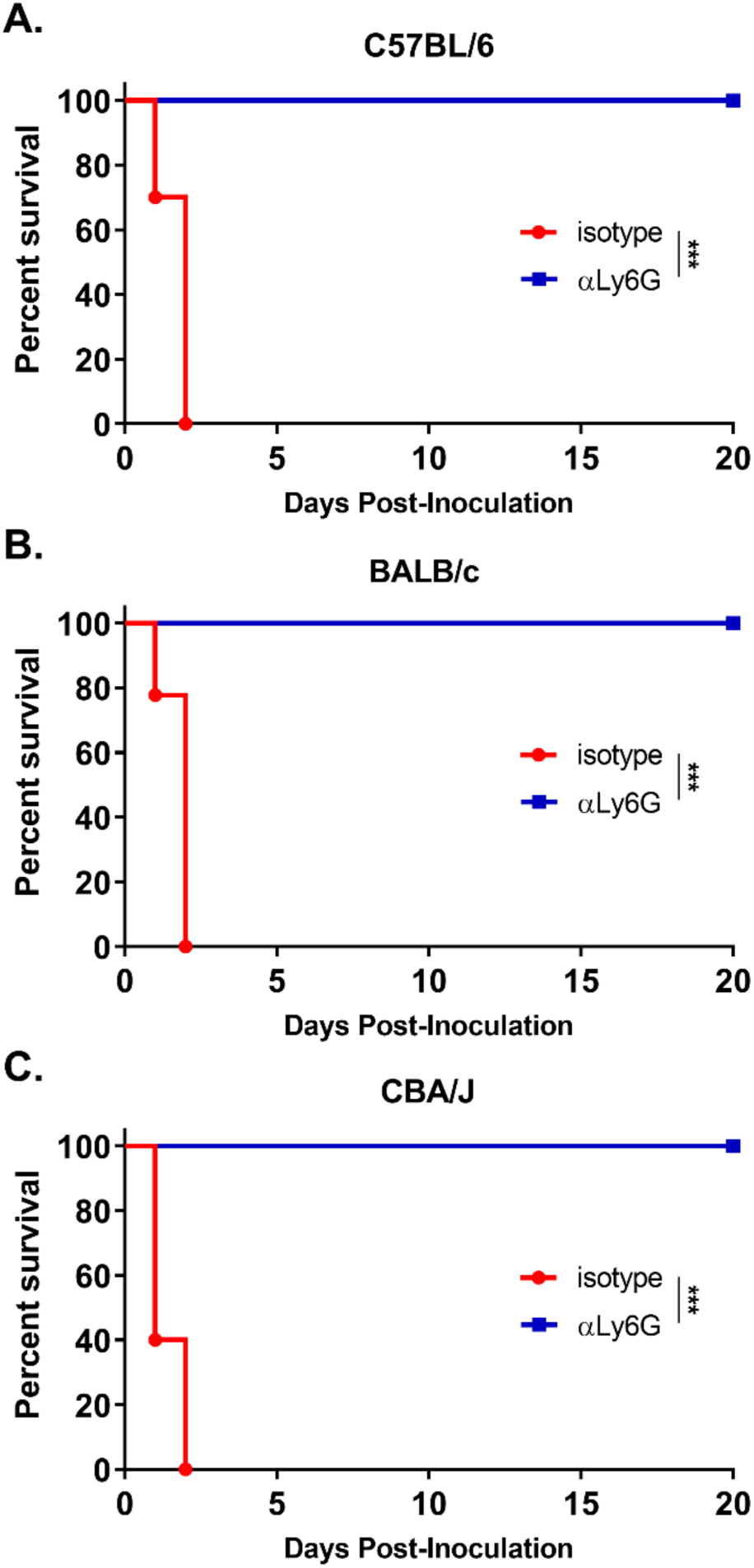
Depletion of neutrophils protects *chs3*Δ inoculated mice: (A) C57BL/6, (B) BALB/c, or (C) CBA/J mice were inoculated with 10^7^ Heat-killed CFUs of each strain by intranasal inoculation. Prior to inoculation and throughout the experiment, mice were treated with isotype antibody or anti-Ly6G antibody. Survival of the animals was recorded as mortality of mice for 20 days post inoculation. Mice that lost 20% of the body weight at the time of inoculation or displayed signs of morbidity were considered ill and sacrificed. Data is cumulative of two independent experiments with 5 mice for *chs3*Δ and *chs3Δ::CHS3* each for a total of 10 mice. Virulence was determined using Mantel-Cox curve comparison with statistical significance determined by log-rank test. (***, *P* < 0.001).

## Discussion

We have previously shown that deletion of a specific chitin synthase (CHS3), or deletion of all three chitin deacetylases causes a significant reduction in chitosan in the vegetative cell wall (9). These chitosan deficient strains of *C. neoformans* were found to be avirulent and rapidly cleared from the murine lung (9). Moreover, infection with a chitosan deficient *C. neoformans* strain lacking three chitin deacetylases (*cda1Δ2Δ3Δ*) was found to confer protective immunity to a subsequent challenge with a virulent wild type counterpart (10). These findings suggest that there is an altered host response to chitosan-deficient strains. Surprisingly, we observed that mice inoculated with chitosan-deficient *chs3*Δ all died within 36 hours (Figs. 2 and 3) and death was associated with an aberrant hyperinflammatory immune response indicating that chitosan is critical in modulating the immune response to *Cryptococcus*.

The immune response to *Cryptococcus*, as well as the magnitude of the response, can play a protective or detrimental role. Our data fits well within the damage-response framework proposed by Casadevall and Pirofski (12) where host damage or benefit is dependent on the host response. This is represented as a parabolic curve, where too little of a response to a microorganism can lead to damage caused by the microorganism and to strong of a host response can lead to damage caused by the host response. This framework is observed in cryptococcal infected AIDS patients. Too little of a response can lead to patient death due to the fungus, whereas a hyperactive response can lead to death caused by immunopathology. AIDS patients treated with antiretroviral therapy often develop cryptococcal immune reconstitution inflammatory syndrome (IRIS) which is an exaggerated and frequently deadly inflammatory reaction that complicates recovery from immunodeficiency (15). Cryptococcal IRIS emphasizes the potential role of the host immune system in mediating host damage and disease symptoms.

There is precedent to study mutants that induce an aberrant hyperinflammatory immune response, as similar responses have been observed with cryptococcal IRIS. Cryptococcal IRIS develops in 8-49% of patients with known cryptococcal disease before antiretroviral therapy (16). The pathogenesis of IRIS is poorly understood, and prediction of IRIS is not currently possible. Innate immune cells, such as monocytes and neutrophils, are of increasing interest in IRIS pathophysiology, since granuloma appears to be frequently found in IRIS lesions (15). Additionally, at the time of IRIS onset multiple proinflammatory cytokine are detected, including IL-6 (17). Further study of the *chs3*Δ immune response could advance our understanding of host immune mechanisms involved in an inappropriately strong immune response to *Cryptococcus*, like those seen in immune reconstitution inflammatory syndrome. These studies have the potential to advance our understanding of a significant problem in the management of cryptococcal patients.

Other cryptococcal mutants that have defects in the cell wall, like *rim101Δ*, have been found to induce a strong proinflammatory response and lead to neutrophil recruitment (18), however, not to the order of magnitude observed with *chs3Δ.* Neutrophils have a complicated role in the cryptococcal immune response. While neutrophils can kill *C. neoformans*, the fungus can modulate the neutrophil response. Cryptococcal capsular and cell wall components can inhibit neutrophil migration (19, 20), and can inhibit the production of neutrophil extracellular traps (21). In the brain, neutrophils have been shown to be important in clearance of the fungus from the microvasculature (22, 23). Neutrophil depletion in a protective immunization model did not affect pulmonary fungal burden, indicating that neutrophils are not required for clearance (13) or for the secondary response (14). These data further support the observation by Mednick et al. that neutropenic mice given a pulmonary *C. neoformans* infection survived significantly longer than control mice that had an intact neutrophil compartment (24), therefore indicating that neutrophils are not necessary for protective responses against cryptococcal infection. We observed a significant increase in neutrophil recruitment to the lungs of mice inoculated with the *chs3*Δ strain (Figs 6B-C). Mice inoculated with HK *chs3*Δ and depletion of neutrophils all survived whereas the isotype treated mice all died (Fig. 7), indicating a detrimental role of neutrophils. Further supporting a harmful role for neutrophils, mice with genetically-induced neutrophilia appear to have increased susceptibility to cryptococcal disease (25). More work is needed to elucidate our understanding the cryptococcus:neutrophil interactions.

In summary, we have shown that inoculation with either live or dead cells from the chitosan deficient strain, *chs3*Δ, leads to death of the mice within 36 hours. The rapid on set of death is likely due to an aberrant hyperinflammatory immune response as mortality was not dependent on viable fungi. Histology, cytokine profiling, and flow cytometry indicates a massive influx of neutrophils in the mice inoculated with *chs3*Δ. Depletion studies show a damaging role for neutrophils in the response to *chs3*Δ. Altogether, chitosan plays a major role in the immune response to *C. neoformans.* In addition, the response to chitosan deficient *C. neoformans* seems to depend on the type of genes deleted, as not all chitosan deficient strains induce the same immune response.

## Materials and Methods

### Fungal strains and media

*C. neoformans* strain KN99*α* was used as the wild-type strain and as progenitor of mutant strains. Strains were grown in YPD broth (1% yeast extract, 2% bacto-peptone, and 2% dextrose) or on YPD solid media containing 2% bacto-agar. Selective YPD media was supplemented with 100 μg/mL nourseothricin (NAT) (Werner BioAgents, Germany).

### Strain construction

Gene-specific deletion construct of the chitin synthase 3 gene (CNAG_05581) was generated using overlap PCR gene technology described previously (26, 27) and included the nourseothricin resistance cassette. The primers used to disrupt the genes are shown in Table S1. The Chs3 deletion cassette contained the nourseothricin resistance cassette resulting in a 1,539 bp replacement of the genomic sequence between regions of primers 3-Chs3 and 6-Chs3 shown in upper case in Table S1. The construct was introduced into the KN99*α* strain using biolistic techniques (28). To generate a *CHS3* complement strain, we replaced the NAT resistance cassette in the *chs3* deletion strain with the native *CHS3* gene sequence by electroporation (29) and screened for NAT sensitivity.

### Morphological analysis

Cells were incubated for 2 days in YPD medium at 30°C with shaking and diluted to an OD_650_ of 0.2 with PBS. Five microliters each cell solution was spotted on to a clean glass slide and photographed on an Olympus BX61 microscope.

### Evaluation of temperature sensitivity

Wild-type, *chs3* deletion and *chs3Δ::CHS3* complement strains were grown in liquid YPD for 2 days at 30°C with shaking. Cells were diluted to OD_650_=1.0 and 10-fold serial dilutions were made. Five microliters of each dilution were spotted on YPD plates and the plates were incubated for 2-3 days at 30°C and 39°C and photographed.

### Cellular chitosan measurement

As previously described, MBTH (3-methyl −2-benzothiazolinone hydrazone) based chemical method was used to determine the chitin and chitosan content (30). In brief, cells were grown in liquid YPD for 2 days at 30°C with shaking collected by centrifugation. Cell pellets were washed two times with PBS, pH 7.4 and lyophilized. The dried samples were resuspended in water first before adding KOH to a final concentration of 6% KOH (w/v). The alkali suspended material was incubated at 80°C for 30 min with vortexing in between to eliminate non-specific MBTH reactive molecules from the cells. Alkali treated material was then washed several times with PBS, pH 7.4 to make sure that the pH of the cell suspension was brought back to neutral pH. Finally, the cell material was resuspended in PBS, pH 7.4 to a concentration of 10mg/mL in PBS (by dry weight) and a 0.1 mL of each samples was used in the MBTH assay (31).

### Mice

BALB/c (000651), CBA/J (000656), and C57BL/6 (000664) mice were obtained from Jackson Laboratory (Bar Harbor, ME). BALB/c and C57BL/6 obtained from Jackson Labs are also known as BALB/cJ and C57BL/6 respectively. All mice were 6 to 8 weeks old at the time of inoculation. All animal protocols were reviewed and approved by the Animal Studies Committee of the Washington University School of Medicine and conducted according to National Institutes of Health guidelines for housing and care of laboratory animals.

### Pulmonary inoculations

Strains were grown at 30°C, 300 rpm for 48 hours in 50 mL YPD. The cells were centrifuged, washed in endotoxin-free 1x PBS and counted with a haemocytometer. For studies utilizing heat-killed organism, after being diluted to the desired cell number in PBS, the inoculum was heated at 70°C for 15 minutes. Complete killing was assayed by platting for CFUs. Mice were anaesthetized with an intraperitoneal injection (200 µL) of ketamine (8 mg/mL)/dexmedetomidine (0.05 mg/mL) mixture and then given an intranasal inoculation with 1 × 10^7^ CFU of live or heat-killed organism in 50 μl of sterile PBS. Anesthesia was reversed by an intraperitoneal injection of (200 µL) of antipamezole (0.25mg/mL). The mice were fed *ad libitum* and monitored daily for symptoms. For survival studies mice were sacrificed when body weight fell below 80% of weight at the time of inoculation. For cytokine analysis, flow cytometry studies, and histology, mice were euthanized at 8-hours post-inoculation by CO_2_ inhalation and the lungs were harvested.

### Histology

Mice were sacrificed according to approved protocols, perfused intracardially with sterile PBS, and the lungs inflated with 10% formalin. Lung tissue was then fixed for 48 hours in 10% formalin and submitted to HistoWiz Inc. (histowiz.com) for histology using a Standard Operating Procedure and fully automated workflow. Samples were processed, embedded in paraffin, sectioned at 4μm and stained using hematoxylin-eosin (H&E). After staining, sections were dehydrated and film coverslipped using a TissueTek-Prisma and Coverslipper (SakuraUSA, Torrance, CA). Whole slide scanning (40x) was performed on an Aperio AT2 (Leica Biosystems, Wetzlar, Germany).

### Cytokine Analysis

Cytokine levels in lung tissues were analyzed using the Bio-Plex Protein Array System (Bio-Rad Laboratories, Hercules, CA). Briefly, lung tissue was excised and homogenized in 2 ml of ice-cold PBS containing 1X Pierce Protease Inhibitor cocktail (Thermo Scientific, Rockford, IL). After a homogenization, Triton X-100 was add to a final concentration of 0.05% and the samples were clarified by centrifugation. Supernatant fractions from the pulmonary homogenates were then assayed using the Bio-Plex Pro Mouse Cytokine 23-Plex (Bio-Rad Laboratories) for the presence of IL-1α, IL-1β, IL-2, IL-3, IL-4, IL-5, IL-6, IL-9, IL-10, IL-12 (p40), IL-12 (p70), IL-13, IL-17A, granulocyte colony stimulating factor (G-CSF), granulocyte monocyte colony stimulating factor (GM-CSF), interferon-γ (IFN-γ), CXCL1/keratinocyte-derived chemokine (KC), CCL2/monocyte chemotactic protein-1 (MCP-1), CCL3/macrophage inflammatory protein-1α (MIP-1α), CCL4/MIP-1β, CCL5/regulated upon activation, normal T cell expressed and secreted (RANTES) and tumor necrosis factor-α (TNF-α).

### Flow Cytometry

Cell populations in the lungs were identified by flow cytometry. Briefly, lungs from individual mice were enzymatically digested at 37°C for 30 min in digestion buffer (RPMI 1640 containing 1 mg/ml of collagenase type IV). The digested tissues were then successively passed through sterile 70 and 40 µm pore nylon strainers (BD Biosciences, San Jose, CA). Erythrocytes in the strained suspension were lysed by incubation in NH_4_Cl buffer (0.859% NH_4_Cl, 0.1% KHCO_3_, 0.0372% Na_2_EDTA; pH 7.4; Sigma-Aldrich) for 3 min on ice, followed by the addition of a 2-fold excess of PBS. The leukocytes were then collected by centrifugation, resuspended in sterile PBS, and were stained using the LIVE/DEAD™ Fixable Blue Dead Cell Stain Kit (1:1000; Invitrogen, Carlsbad, CA) for 30 min at 4° C in the dark. Following incubation, samples were washed and resuspended FACS buffer (PBS, 0.1% BSA, 0.02%NaN_3_, 2mM EDTA) and incubated with CD16/CD32 (Fc Block™; BD Biosciences, San Jose, CA) for 5 min. For flow cytometry, 1×10^6^ cells were incubated for 30 min at 4° C in the dark with optimal concentrations of fluorochrome-conjugated antibodies (Table S2 for antigen, clone and source) diluted in Brilliant Stain Buffer (BD Biosciences). After three washes with FACS buffer, the cells were fixed in 2% ultrapure paraformaldehyde. For data acquisition, >200,000 events were collected on a BD LSRFortessa X-20 flow cytometer (BD Biosciences, San Jose, CA), and the data were analyzed with FlowJo V10 (TreeStar, Ashland, OR). The absolute number of cells in each leukocyte subset was determined by multiplying the absolute number of CD45^+^ cells by the percentage of cells stained by fluorochrome-labeled antibodies for each cell population analyzed.

### Neutrophil depletion

Mice were depleted of neutrophils via intraperitoneal (ip) administration of 200 μg anti-Ly6G (clone 1A8; BioXcell) in 100 μl. Control mice received 200 μg IgG2a isotype control antibody (clone 2A3; BioXcell) in 100 μl. Depletions were started 24 hours prior to challenge and the mice were injected every 24 hours for the first 5 days post challenge. After day 5, the mice were injected every 48 hours.

### Statistics

Data were analyzed using GraphPad Prism, version 8.0 (GraphPad Software, Inc., La Jolla, CA). The one-way analysis of variance (ANOVA) with the Tukey’s multiple-correction test was used to compare more than two groups. Kaplan-Meier survival curves were compared using the Mantel-Cox log rank test. p values <0.05 were considered significant.

## Acknowledgments

This work was funded by National Institutes of Health grants AI072195 and AI125045 to JKL. CRH was partly funded by a National Institute of Allergy and Infectious Diseases training grant (T32 AI007172). The funders had no role in study design, data collection and interpretation, or the decision to submit the work for publication.

---

**Supplemental figure 1:**
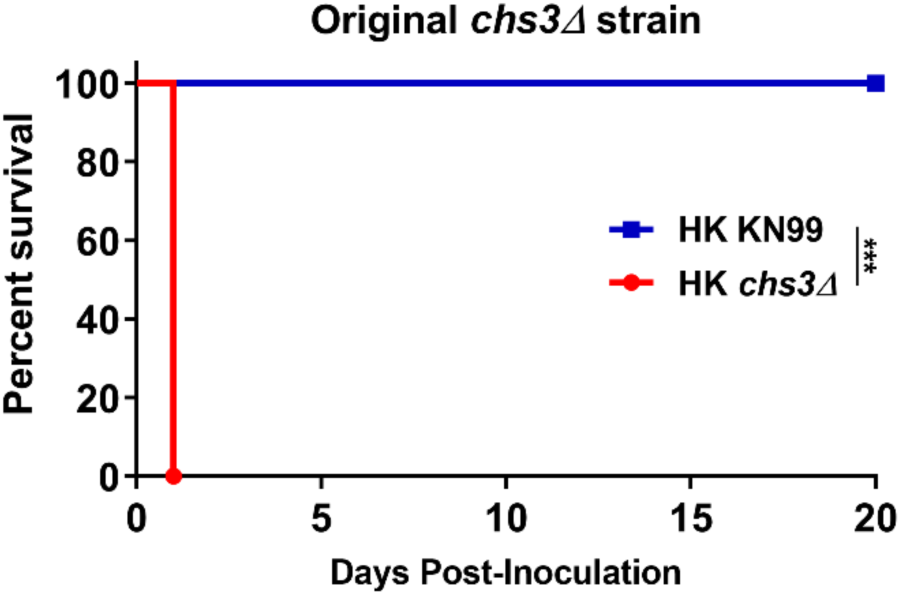
The original *chs3*Δ strain also induces rapid mortality: C57BL/6 mice were inoculated with 10^7^ Heat-killed CFUs of each strain by intranasal inoculation. Survival of the animals was recorded as mortality of mice for 20 days post inoculation. Mice that lost 20% of the body weight at the time of inoculation or displayed signs of morbidity were considered ill and sacrificed. Data is representative of one experiment with 5 mice for each strain. Virulence was determined using Mantel-Cox curve comparison with statistical significance determined by log-rank test. (***, *P* < 0.001).

**Supplemental figure 2.**
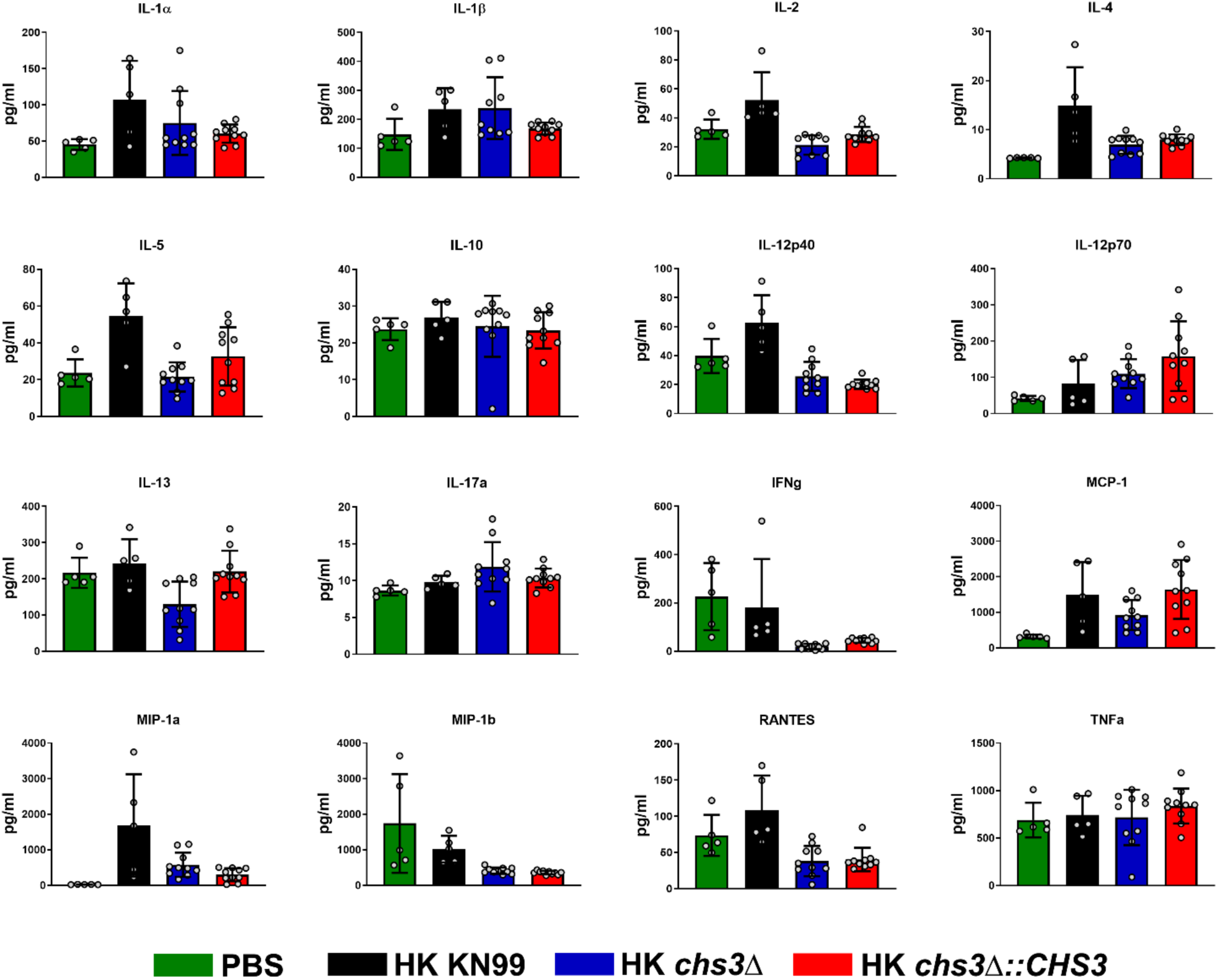
Cytokine/chemokine analysis: C57BL/6 mice were inoculated with 10^7^ Heat-killed CFUs of each strain by intranasal inoculation. At 8 hours post inoculation, homogenates were prepared from the lungs of each group. Cytokine/chemokine responses were determined from the lung homogenates using the Bio-Plex Protein Array System. Data is cumulative of one experiment with 5 mice for PBS and KN99, and two experiments with 5 mice for *chs3*Δ and *chs3Δ::CHS3* each for a total of 10 mice experiments, ± standard errors of the means (SEM). Each dot represents data from an individual mouse.

**Supplemental figure 3.**
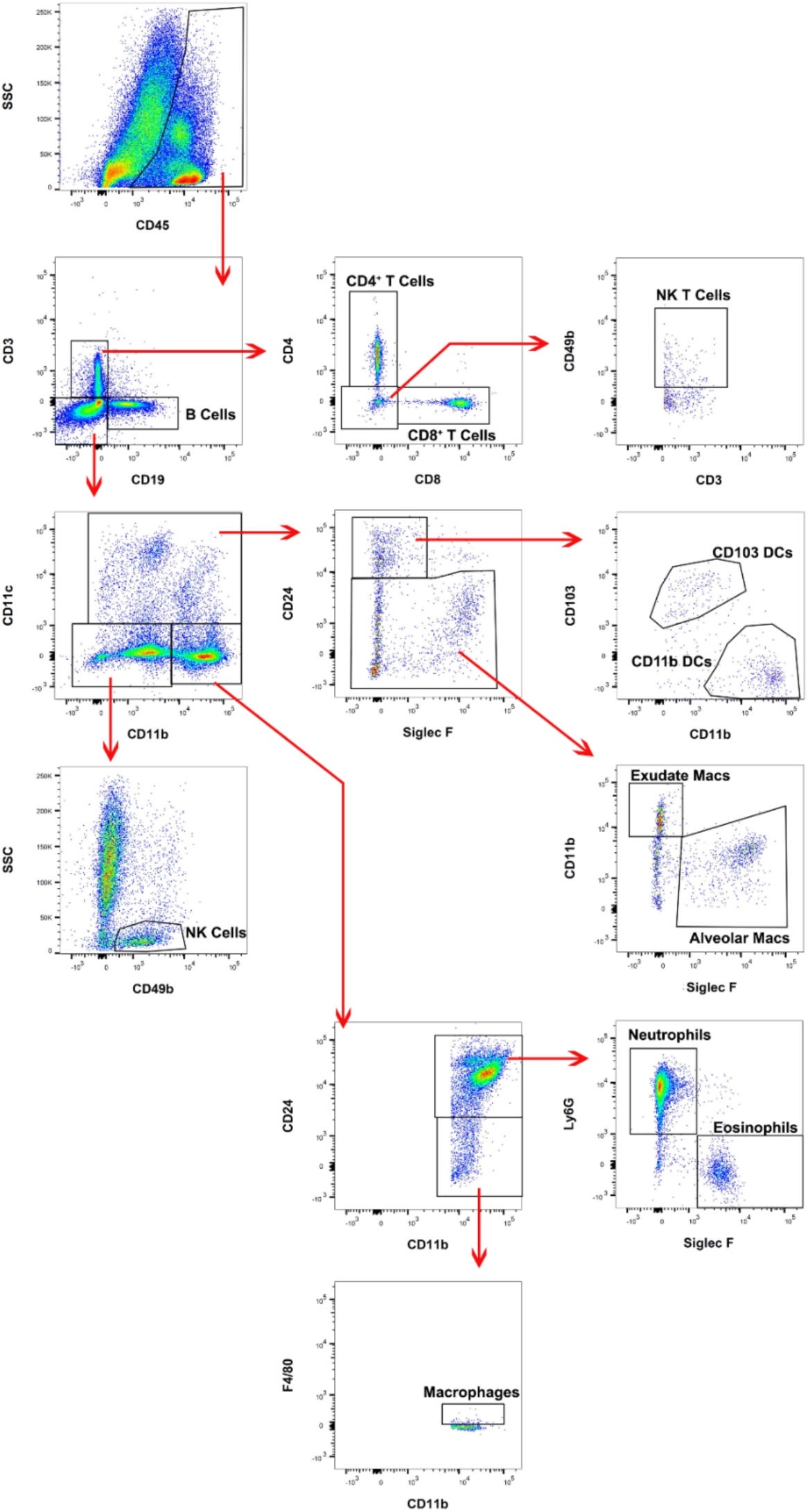
Flow cytometry gating strategy. Flow cytometry analysis of host leukocyte populations in C57BL/6 mice 8 hours post inoculation with 10^7^ Heat-killed CFUs of *chs3*Δ. After separation based on physical properties to eliminate debris and doublets, dead cells were excluded by live/dead staining (not shown). CD45+ cells (leukocytes; top left) were subjected to analysis based on their expression of CD3, CD4, CD8a, CD11b, CD11c, CD19, CD24, CD49b, CD103, F4/80, Ly-6G, and Siglec F. The specific populations of cells identified are indicated on the flow plots.

**Supplemental figure 4.**
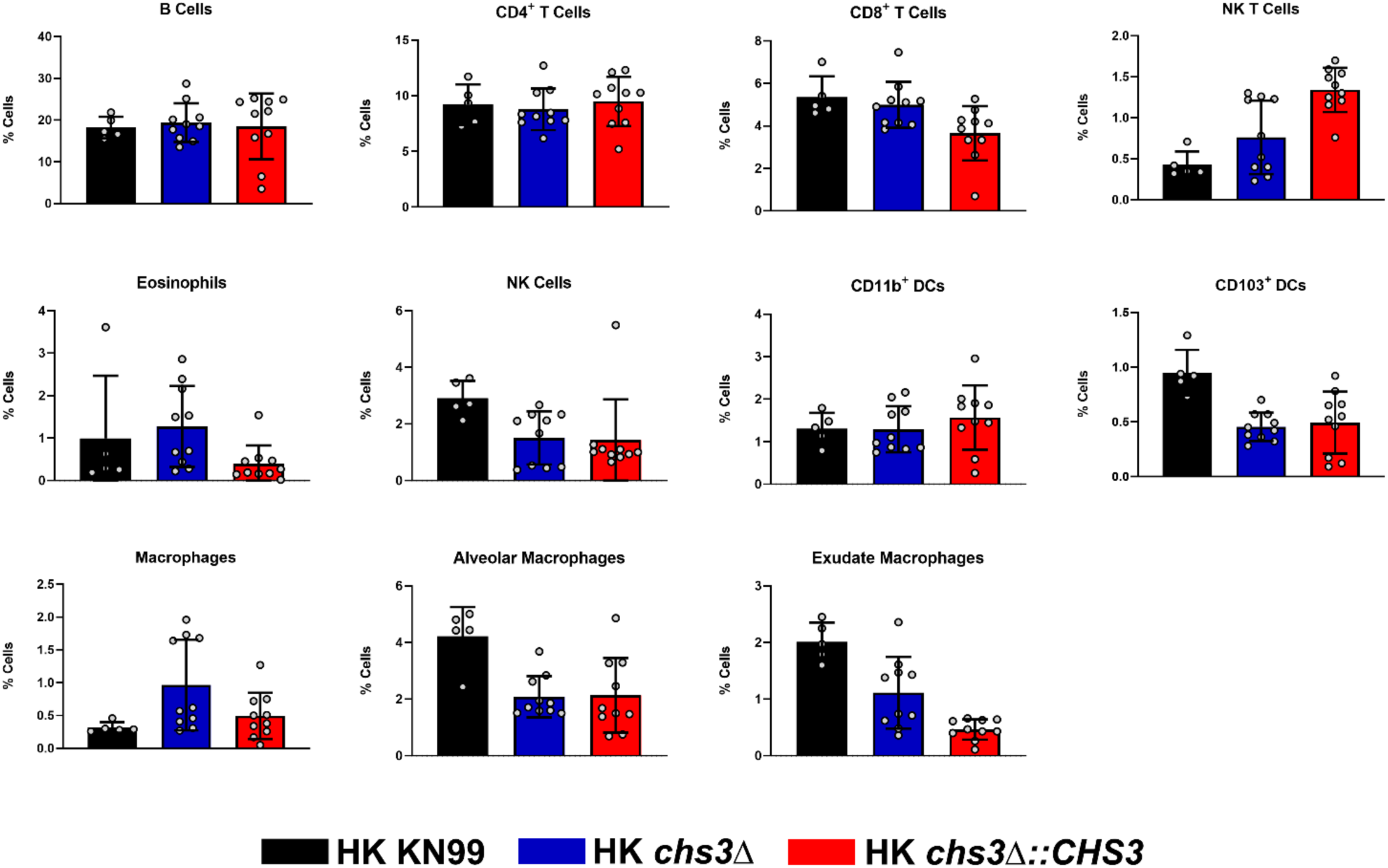
Flow cytometry analysis: C57BL/6 mice were inoculated with 10^7^ Heat-killed CFUs of each strain by intranasal inoculation. At 8 hours post inoculation, pulmonary leukocytes were isolated from the lungs of each group and subjected to flow cytometry analysis Data is cumulative of one experiment with 5 mice for PBS and KN99, and two experiments with 5 mice for *chs3*Δ and *chs3Δ::CHS3* each for a total of 10 mice experiments, ± standard errors of the means (SEM). Each dot represents data from an individual mouse.

**Table S1.**
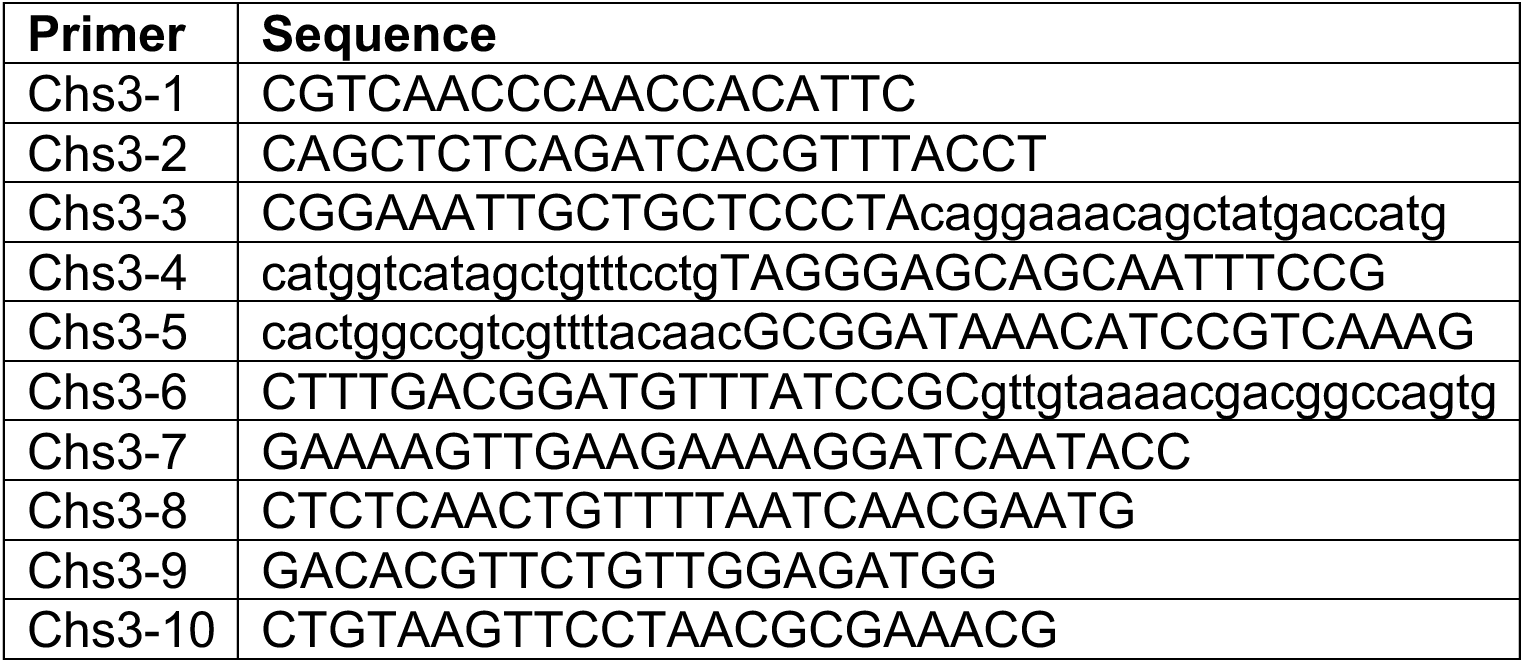
Primer used in this study.

**Table S2.** Antibodies for flow analysis.

| Antigen | Clone | Fluorophore | Dilution | Company |
| --- | --- | --- | --- | --- |
| CD3 | 17-A2 | APC-Cy7 | 1:250 | BioLegend |
| CD4 | GK1.5 | BUV737 | 1:500 | BD Biosciences |
| CD8a | 53-6.7 | BV650 | 1:125 | BD Biosciences |
| CD11b | M1/70 | BV510 | 1:250 | BD Biosciences |
| CD11c | N418 | PE | 1:125 | Invitrogen |
| CD16/CD32 Fc Block | 2.4G2 | (not applicable) | 1:500 | BD Biosciences |
| CD19 | 1D3 | BV786 | 1:125 | BD Biosciences |
| CD24 | M1/69 | PerCP-Cy5.5 | 1:125 | BD Biosciences |
| CD45 | 30-F11 | Pacific Blue | 1:250 | BioLegend |
| CD49b | DX5 | PE-Cy7 | 1:250 | Invitrogen |
| CD103 | 2E7 | Alexa488 | 1:125 | BioLegend |
| F4/80 | BM8 | APC | 1:250 | Invitrogen |
| Ly6G | 1A8 | BV711 | 1:125 | BD Biosciences |
| Siglec-F | E50-2440 | PE-CF594 | 1:250 | BD Biosciences |

